# Dolphin social phenotypes vary in response to food availability but not the North Atlantic Oscillation index

**DOI:** 10.1101/2023.04.04.535551

**Authors:** David N. Fisher, Barbara J. Cheney

## Abstract

Social behaviours can allow individuals to flexibly respond to environmental change, potentially buffering adverse effects. However, individuals may respond differently to the same environmental stimulus, complicating predictions for population-level response to environmental change. Here we show that bottlenose dolphins (*Tursiops truncatus*) alter their social behaviour at yearly and monthly scales in response to a proxy for food availability (salmon abundance) but do not respond to variation in a proxy for climate (the North Atlantic Oscillation index). There was also individual variation in plasticity for gregariousness and connectedness to distant parts of the social network, although these traits showed limited repeatability. In contrast, individuals showed consistent differences in clustering with their immediate social environment at the yearly scale but no individual variation in plasticity for this trait at either time scale. These results indicate that social behaviour in free-ranging cetaceans can be highly resource dependent with individuals increasing their connectedness over short timescales but possibly reducing their wider range of connection at longer timescales. Some social traits showed more individual variation in plasticity or mean behaviour than others, highlighting how predictions for the responses of populations to environmental variation must consider the type of individual variation present in the population.

## Introduction

Animals engage in social interactions with conspecifics which are fundamental for determining health, access to resources, and reproductive success [1]. Consequently, social interactions have a strong influence on ecological processes such as population dynamics and evolutionary processes such as the response to selection [2–4]. To maintain the best fit with their environment, animals may adjust their social behaviour as conditions change [5], for instance being more gregarious when resources are plentiful but less tolerant of conspecifics when resources are scarce [6]. Animals may also change their behaviour through development and during senescence [7], and may non-adaptively adjust their behaviour due to direct effects of the environment and other limitations [8]. Such plasticity is a hallmark of behavioural traits and gives behaviour an important role in how animals interact with their environment.

When trying to understand how animals may respond plastically to changing environments, most examine responses at the population level [e.g., 9], presuming that any individual variation in response is absent or simply aggregates to give the population-level response. However, individuals may show variation in plasticity, and so each will respond differently to change [10,11, sometimes referred to as ‘I x E’ i.e., individual by environment interactions,12,13]. For example, European field crickets (*Gryllus campestris*) become bolder and more active as they age, but individuals vary in the extent of this, with some not increasing or even decreasing [14]. The degree of plasticity animals show can be correlated with their mean behaviour (an “intercept-slope correlation”) which determines how the magnitude of among-individual differences varies across environments and indicates the extent to which plasticity is a separate trait in its own right [11,15]. Individuality in plasticity can influence biological processes such as population growth and adaptive change at a range of scales [16, see: 17 and accompanying papers], giving fundamentally different results to when population-level only effects are assumed [15,18]. For example, Seebacher and Little demonstrated that mosquitofish (*Gambusia holbrooki*) differ in how their performance changes with temperature, resulting in a switching of the rank of swimming speed of individuals between cool and warm temperatures. This changed which individuals might be predated, with a lower fraction of the population reaching the critical speed to avoid predation in cool temperatures [19]. Therefore, variation in plasticity will alter both the strength of selection and which genotypes produce phenotypes that are selected for, altering evolution trajectories. Additionally, the extent of among-individual variation in plasticity gives an upper-limit to the heritability of plasticity, which indicates how rapidly plasticity itself can evolve [20–22]. Understanding how plasticity as a trait in its own right can evolve is key for understanding how animals will adapt to more variable climates [22,23].

Marine mammals are a key group to study individual variation in response to environmental change. They are typically long-lived, increasing the relative importance of plasticity versus adaptive evolution for coping with contemporary environmental change [24]. They are also exposed to a wide range of changing conditions during their lifetimes including climate, food availability, and pollution, and their populations are often of conservation concern. All of these factors increase the need for us to understand how they respond to changes in their environment [25–32].

Here, we studied a population of bottlenose dolphins (*Tursiops truncatus*) in the North Sea for over 30 years, regularly recording their social associations. Previous work in this study population has demonstrated that critical group sizes increase in years of higher salmon abundance [33], and we extend this by examining multiple facets of individual-level social behaviour at both the monthly and yearly temporal scales. We achieved this by using social network analysis to quantify three different dimensions of individual social behaviour: an individual’s gregariousness (strength), how tightly its immediate social group interact together (clustering coefficient), and how well connected an individual is to the entire population (closeness; each described in more detail below). We then used random regression models to quantify individual social phenotypes and determine how these social phenotypes depend on yearly and monthly variation in available proxies for climate (at a broad scale) and food availability (at a local scale). Our analyses also indicated whether bottlenose dolphins show individual changes in response to the environment, or if population-level change was more prominent.

Specifically, we were interested in how social behaviour depends on current environmental conditions. Social behaviours are often highly dependent on both resource availability and spatial distribution [6] and current climatic conditions can impose energetic constraints on individuals [34,35] and impact their ability to move around their environment [36]. Bottlenose dolphin group sizes off the north west coast of Spain showed a nonlinear relationship with the North Atlantic Oscillation (NAO) index [37], while abundances of Indo-Pacific bottlenose dolphins (*T. aduncus*) are impacted by a combination of the El Nino Southern Oscillation and season [38]. In our study system, previous work indicated that the NAO index at a two-year lag was associated with dolphin critical group size, but this appears to be entirely mediated through food availability [33], something we are testing for directly. As such, we did not consider lagged effects here. We summarised climate through the NAO index (see Methods), where positive values in this region indicate warmer and wetter periods which would make rougher sea conditions, potentially resulting in the dolphins travelling shorter distances. We therefore expect higher NAO values to lead to higher clustering coefficients and lower closeness, but not to affect strength, at monthly and yearly scales. We summarised resource availability through salmon abundances (see Methods). We expect that higher salmon abundances allow dolphins to form larger groups (as found previously) and travel shorter distances to find sufficient food, leading to higher strengths, higher clustering coefficients, and lower closeness at both temporal scales.

## Material and methods

### Study site & group data collection

This study used data from a bottlenose dolphin population on the east coast of Scotland (Fig. 1). The population of over 200 individuals [39] has been studied intensively as part of a long-term individual based study [40–42]. We use data from boat-based photo-identification surveys carried out annually between 1990 and 2021 which regularly recorded dolphin groups within the Moray Firth Special Area of Conservation (SAC; 92/43/EEC), a core part of the population’s range which over 50% of the population use each year (40). All surveys were made from small (5-6m) boats with outboard engines, carefully and slowly manoeuvring the boat around each group to obtain high quality images of the left and right side of as many dorsal fins as possible. Surveys initially followed a fixed survey route until 2001 when, as a result of changing dolphin distribution within the SAC, flexible survey routes were introduced to maximise sightings probability [more details in:, 43]. Data were available from a total of 690 surveys (between 9 and 35 surveys each year; average of 22) with the majority carried out between May to September.

**Figure 1.**
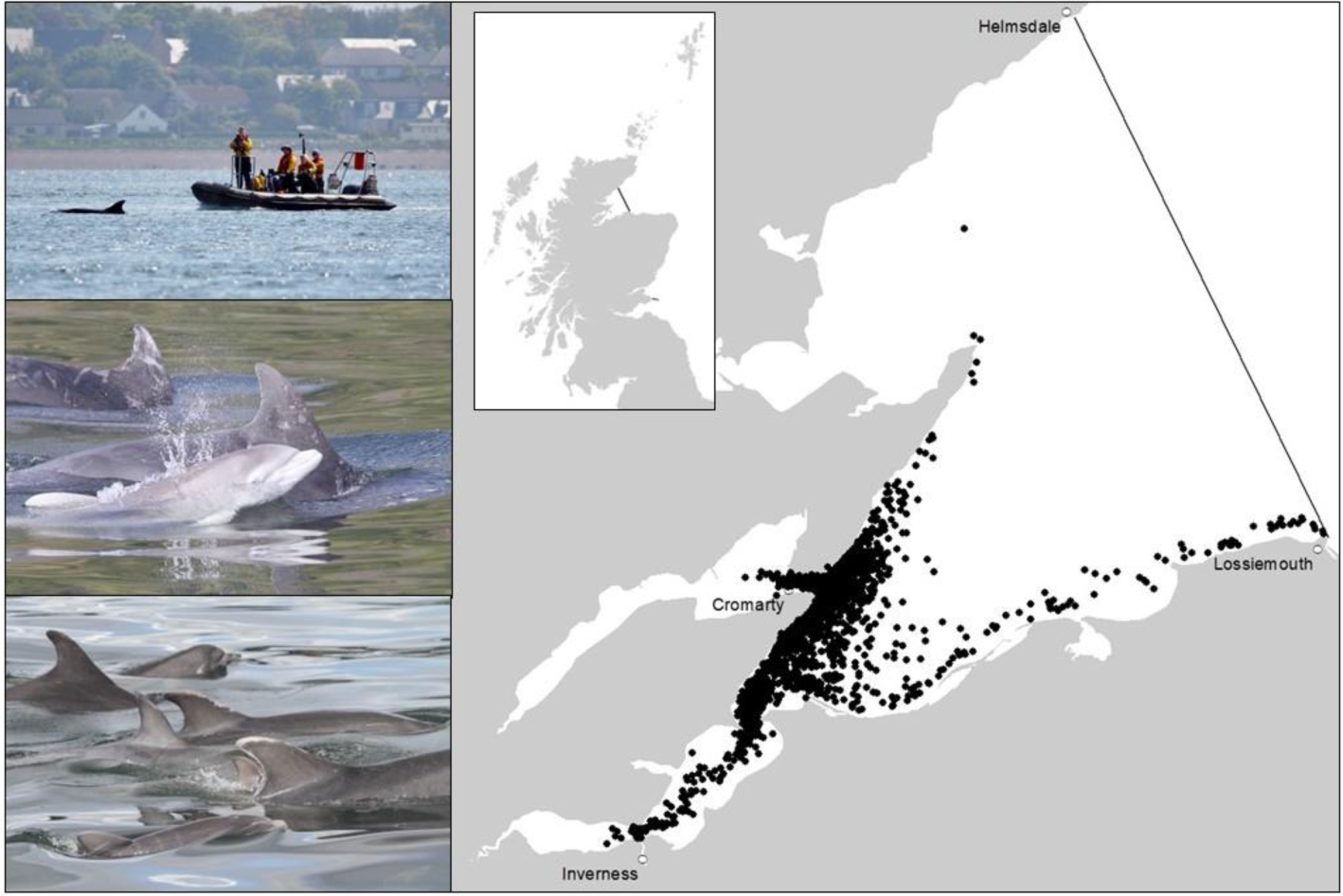
Images depicting a photo-identification survey (top left), a newborn calf with mother (middle left), unique markings used to identify individuals (bottom left), and location of encounters with bottlenose dolphins within the Moray Firth Special Area of Conservation from 1990 to 2021 (right).

During surveys, when we located a bottlenose dolphin group (one or more individuals in close proximity within 100m, hereafter an “encounter”) we collected photo-identification data following a standardised protocol [43]. We identified individuals from high-quality photographs based on unique markings matched against a photo catalogue of previously identified individuals from the area [40,41,44]. On average in a group 84% are successfully photographed, a rate of identification well above the level at which social network metrics in incomplete networks are reliable [45]. All individual identifications from photographs were confirmed by at least two experienced researchers. For individuals first sighted as calves we could determine their year of birth and so their age [46], but for individuals first sighted as juveniles or adults their exact age is unknown. Sex was determined using genital photographs or if an adult was seen in repeat associations with a known calf [46].

### Social network construction

Individuals sighted during the same encounter were assumed to be in the same group and therefore associating [known as the ‘Gambit of the Group’ 47]. Aggregating many of these records of groups allows one to infer which individuals are frequently associated and which individuals infrequently or never associate. We removed observations of individuals younger than three years old (n = 2668 observations of 242 individuals), as these individuals are not likely to be independent of the mother and so their social associations most likely represent her preferences. We then converted the records of encounters into group by individual matrices (indicating which individuals were seen together in each encounter) and then into weighted, undirected social networks using the R package *asnipe* [48]. Edge weights were set as the simple ratio index, where the number of times two individuals are seen together is divided by the total number of times they are seen, both together and apart [49]. This measure ranges from 0 (individuals never seen together) to 1 (individuals always seen together). We did this separately for each year, creating yearly social networks to assess how social phenotypes vary at this temporal scale in response to environmental conditions. To assess how social phenotypes vary at the monthly scale in response to environmental conditions we then reconstructed social networks per month and removed any months with fewer than 10 encounters [excluding 110 out of 215 months; as networks constructed using fewer than 10 observations can be biased;, 50]. Histograms of the frequency of the number of encounters per year and per month are shown in Fig. S1.

For each individual present in the network, for each year and again for each month we calculated three network metrics. First was “strength”, the sum of all an individual’s associations, which as our associations are based on observations of co-occurrence in groups is analogous to typical size of groups an individual is in. Second was “weighted clustering coefficient”, the rate at which an individual associates with other individuals who also associate with each other. This metric represents how tightly individuals interact in their immediate social environment (possibly analogous to “alliances” between three or more individuals, see also: [51,52]), at the expense of interacting with a wider range of individuals. Finally, we quantified “closeness”, the inverse of the mean of the path lengths between that individual and each other individual in the network, corrected for network size to allow comparisons among networks which vary in the number of individuals. Closeness represents the dolphin’s connectedness to the wider population, and would be high if an individual linked two communities or moved between different areas each containing more sedentary individuals.

We removed an individual’s scores for a given year if they had fewer than five observations in that year (removing 811 observations and leaving 874), as the social network position of those individuals would be highly uncertain. They would however still contribute to the social environments and therefore social network measures of individuals in that year who had five or more observations. We repeated this at the month level, removing individuals’ monthly scores when they had fewer than five observations that month (removing 3423 observations and leaving 320). Histograms of the frequency of the number of encounters per individual per year and per month are shown in Fig. S2. We had initially performed the analysis with a threshold of three observations before switching to the higher threshold of five; see the supplementary materials for the results with the lower threshold. The social network measures were not strongly correlated; Pearson correlations between individuals’ strength and clustering coefficient were 0.141 (yearly) and 0.346 (monthly), for strength and closeness they were −0.004 (yearly) and −0.245 (monthly), and for clustering coefficient and closeness they were 0.129 (yearly) and 0.170 (monthly).

### Environmental data

We used the NAO index in the same time period the grouping observations were made as a measure of climate. We used monthly and yearly measures of the NAO index between 1990 and 2021 downloaded from https://www.cpc.ncep.noaa.gov/products/precip/CWlink/pna/nao.shtml (Fig. S3). This index indicates the atmospheric pressure difference between the low pressure zone over Iceland and the high pressure zone over the Azores [53,54]. This index has frequently been linked to the ecology of animal populations [55,56], for example influencing the foraging behaviour of Cory’s shearwaters [*Calonectris borealis*;, 57]. Climatic effects on cetaceans are typically thought to occur via changes in prey species [31,58]; for instance Lusseau *et al*. found the NAO at a two-year lag influenced critical group size in our study population through the lagged variable’s effect on food availability [33]. However, it is also possible that cetaceans respond directly to climate, sometimes at even faster rates than their prey species [37,38,59].

For our index of food availability we followed Lusseau *et al*. [33], using data on monthly catches by fishing rods (as opposed to nets) from the wild of both one season and multiple season adult Atlantic salmon (*Salmo salar*) from the Alness, Beauly, Canon, Ness, and Nairn rivers. These feed into the sea where observations of dolphin groups take place and hence are expected to be a good proxy for salmon availability in that area. Further, catches on rods are positively correlated among rivers and among months and with automatic counter data, indicating they are a good proxy for actual abundance [60,61]. Atlantic salmon are an important food source for this dolphin population [62,63], with dolphins forming larger groups when salmon are more abundant [33]. We downloaded monthly data from https://marine.gov.scot/data/marine-scotland-salmon-and-sea-trout-catches-salmon-district-shinyapp (using “rod data” and summing retained and released fish for both MSW and 1SW) and summed monthly catches within a calendar year for yearly measures of fish abundance (Fig. S4).

### Data analysis

All analyses were performed in R [ver 4.3.1;, 64] using linear mixed-effect models in *glmmTMB* [65]. Using regression-based models as opposed to randomisation-based tests has been recommended for analysing questions about node-level social network traits as it improves the ability to make inference while accounting equally well as node permutations for common types of data non-independence [66]. We fitted 12 models, with all combinations across the three social traits, the two environmental variables, and the yearly and monthly timescales. Clustering coefficient cannot be calculated when an individual only associates with one other individual, and so the datasets for these models were slightly smaller than the dataset for strength and closeness (see below). We included individuals with unknown birth and death dates to maximise our sample size, and so neither age nor lifespan could be included as predictor variables. In all models we included the fixed effect of sex [as males and females can differ in social behaviour;, 67,68; ranging behaviour, and survival;, 69], and either the NAO index or the count of caught salmon for that month or year. We did not include both the NAO index and salmon abundance in the same model as we encountered estimation problems with two random slopes. Including sex meant we excluded individuals of unknown sex (140 observations of 66 individuals for the yearly networks, 30 observations of 21 individuals for the monthly networks), but the interactions between individuals of known and unknown sex were still used to build the networks and so associations with individuals of unknown sex still influenced the social network traits of males and females. We mean centred and scaled to unit variance the environmental variable [70], and included the interaction between it and sex, to see if male and female social behaviour responded differently to environmental variation. Random effects were the random intercept for individual ID, the random slope of individual ID with the environmental variable fitted as a fixed effect in the model (again mean centred and with a standard deviation of 1), and the correlation between these two terms. We also included a temporal autocorrelation term (ar1) among years to account for unmodelled environmental variation that changes slowly across years, which could influence social behaviour to make adjacent years more similar than non-adjacent years. Similarly, in the models for monthly variation we included a random effect of month alongside the yearly temporal autocorrelation term. We used a Gaussian error structure for all models. For strength and closeness, we used log link functions as the distributions were right skewed. For clustering coefficient, which is bounded between 0 and 1, we used a logit link function, which is preferable to an arcsine transformation when handling response variables bounded in this way [71]. We used the default optimising algorithm for all models except for both the yearly models for clustering coefficient and closeness, the model for strength in response to monthly variation in the NAO index, and the models for clustering coefficient and closeness in response to monthly variation in salmon abundance, where we used the “BFGS” optimiser, as otherwise the models did not converge [65].

We report the coefficients and standard errors for each fixed effect, along with p-values, to give an idea of the magnitude and uncertainty of each effect. We used the p-values from the Anova function of the *car* package [72], using a Chi-squared test with type III sum of squares. We describe these p-values in terms of “clarity” rather than “significance”; see Dushoff *et al*. [73] for a discussion on this. To test whether individuals clearly differed in their response to the environmental variable, we first tested whether there was a correlation between an individual’s plasticity and its mean behaviour by re-fitting each model (12 in total) with the correlation between random intercepts and slopes suppressed to zero and conducted a likelihood ratio test between the full model and this reduced model with a single degree of freedom. If there was a clear difference between the models, we concluded that the correlation between intercepts and slopes was non-zero. If there was no clear difference between the models, we then tested the importance of the random slopes by comparing the model with an intercept-slope correlation of zero to a model without the random slopes (but still with the random intercepts) using a likelihood ratio test with a mix of zero and one degree of freedom [as is appropriate for testing the clarity of single variance components, 74]. If there was a clear difference between the models, we concluded that individuals differ in their response to the environmental variable. When there is variation in plasticity, the magnitude of among-individual differences varies across environments. We therefore calculated the marginal repeatability for each trait following Schielzeth and Nakagawa [75]. We did this for all trait-environmental variable-time scale combinations (12 in total), even when there was no evidence for random slopes, to aid comparison among traits. Data and R code are available online [76].

## Results

### Change with environmental variables at a yearly scale

For the analysis of how dolphin social phenotypes change at the yearly scale in response to environmental variables, our dataset included 129 unique individuals. For strength and closeness there were 874 measures, and for clustering coefficient there were 873 measures. All traits had a mean of 6.78 (sd = 5.39) measures per individual.

Dolphins’ strength and clustering coefficient were not affected by the NAO index (Fig. 2a and b) or salmon abundance (Fig. 2d and e) in both sexes, and the sexes did not differ in mean strength or clustering coefficient (Table 1, full model results in Tables S1-4, Supplementary Materials). For both strength and clustering coefficient with the NAO index there were no intercept-slope correlations and the random slopes were not statistically clear (Table 2). Individuals did differ in how their strengths changed with salmon abundance, with a clear negative intercept-slope correlation (Table 2). There was no intercept-slope correlation and no random slopes for the change of clustering coefficient with salmon abundance (Table 2). The marginal repeatabilities for strength were low: 0.017 in the NAO index model and 0.018 in the salmon abundance model, suggesting the trait has limited repeatability. The marginal repeatabilities for clustering coefficient were higher: 0.172 and 0.178 for the NAO index and salmon abundance models respectively.

**Figure 2.**
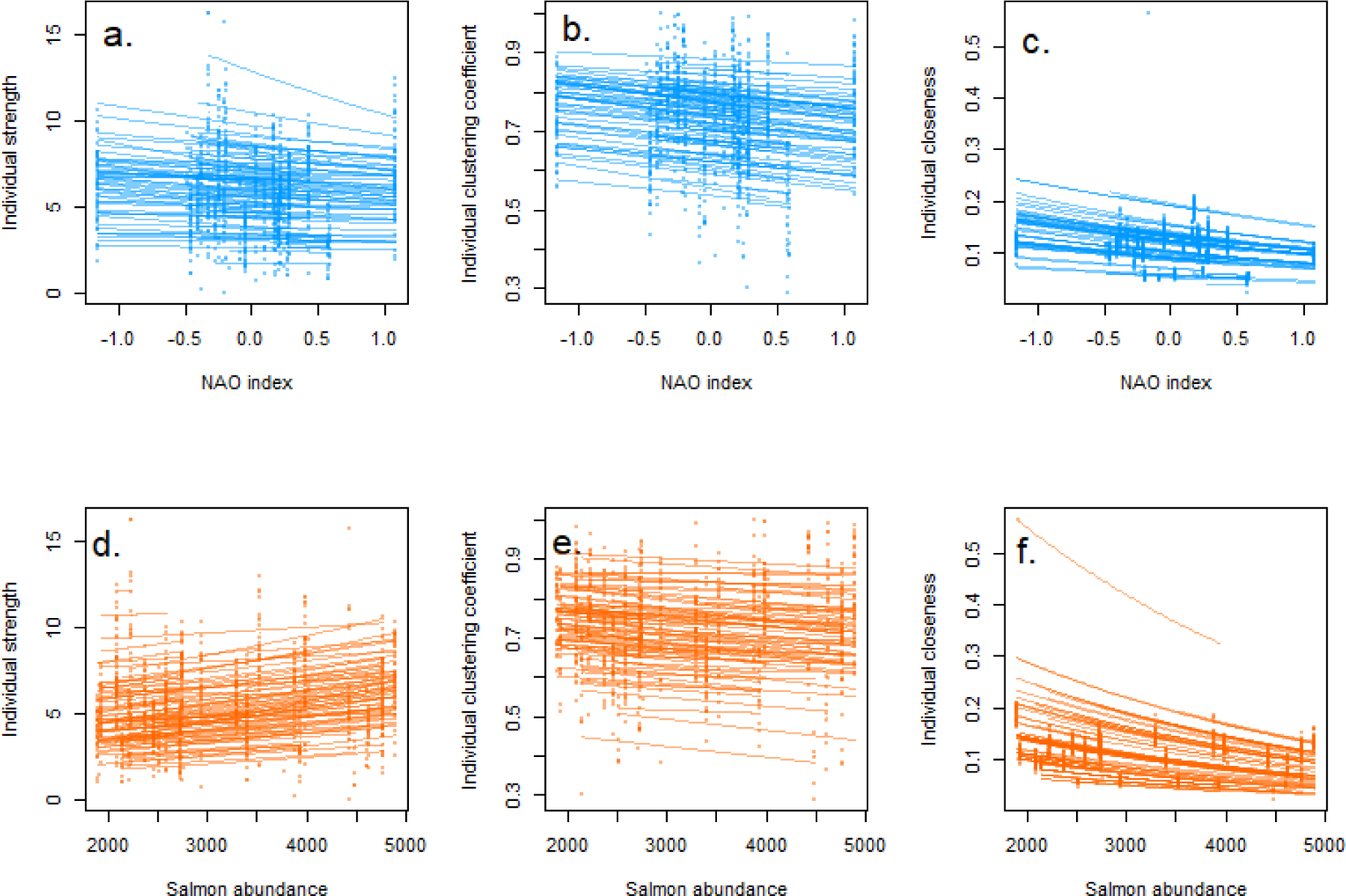
Plots of each of the three social network traits and yearly variation in the North Atlantic Oscillation (NAO) index (a. strength, b. clustering coefficient, c. closeness) and salmon abundances (d. strength, e. clustering coefficient, f. closeness). For each individual dolphin we have predicted its network trait on the observed scale based on the model results, using the “predict” function in R with the individual’s sex, the range of NAO values or salmon counts that individual was exposed to, and picking a random year that individual experienced.

**Table 1.** Main effects (“M”) and interactions with sex (“I”, also shaded in grey) for the two environmental variables effects on the three social network traits, at each of the monthly and yearly scales. Effects clearly different from zero (p < 0.05) are highlighted in bold, while effects with p values between 0.07 and 0.05 are shown in italics.

|  |  |  | Strength | Clustering coefficient | Closeness |
| --- | --- | --- | --- | --- | --- |
| NAO index | Yearly | M | $\beta = -0.035 \pm 0.066, \chi^2 = 0.280, p = 0.597$ | $\beta = -0.102 \pm 0.081, \chi^2 = 1.595, p = 0.207$ | $\beta = -0.115 \pm 0.102, \chi^2 = 1.275, p = 0.259$ |
| | | I | $\beta = -0.002 \pm 0.021, \chi^2 = 0.005, p = 0.942$ | $\beta = 0.031 \pm 0.035, \chi^2 = 0.793, p = 0.373$ | $\beta = 0.006 \pm 0.008, \chi^2 = 0.458, p = 0.499$ |
| | Monthly | M | $\beta = 0.053 \pm 0.034, \chi^2 = 2.437, p = 0.119$ | $\beta = -0.102 \pm 0.056, \chi^2 = 3.304, p = 0.069$ | $\beta = 0.023 \pm 0.033, \chi^2 = 0.464, p = 0.496$ |
| | | I | $\beta = 0.039 \pm 0.037, \chi^2 = 1.105, p = 0.293$ | $\beta = 0.057 \pm 0.069, \chi^2 = 0.678, p = 0.410$ | $\beta = 0.031 \pm 0.037, \chi^2 = 0.713, p = 0.398$ |
| Salmon | Yearly | M | $\beta = 0.108 \pm 0.069, \chi^2 = 2.422, p = 0.120$ | $\beta = -0.027 \pm 0.091, \chi^2 = 1.595, p = 0.207$ | **$\beta = -0.258 \pm 0.094, \chi^2 = 7.489, p = 0.006$** |
| | | I | $\beta = -0.009 \pm 0.026, \chi^2 = 0.126, p = 0.723$ | $\beta = -0.070 \pm 0.038, \chi^2 = 0.794, p = 0.373$ | $\beta = -0.002 \pm 0.008, \chi^2 = 0.091, p = 0.763$ |
| | Monthly | M | **$\beta = 0.137 \pm 0.038, \chi^2 = 12.755, p < 0.001$** | $\beta = 0.116 \pm 0.062, \chi^2 = 3.569, p = 0.059$ | **$\beta = 0.087 \pm 0.040, \chi^2 = 4.714, p = 0.030$** |
| | | I | $\beta = -0.011 \pm 0.045, \chi^2 = 0.065, p = 0.799$ | $\beta = -0.105 \pm 0.074, \chi^2 = 2.011, p = 0.156$ | $\beta = 0.009 \pm 0.030, \chi^2 = 0.081, p = 0.777$ |

**Table 2.** Intercept slope correlations (“ISC”) and their statistical tests, and whether random slopes (“RS”) were present or absent and their statistical tests (likelihood ratio tests in all cases, an asterix is given for the test of the random slopes if it was not performed as the intercept-slope correlation was first found to be clear). Clear positive correlations are noted in bold text and highlighted with blue fill, and clear negative correlations with bold text and orange fill, while clear random slopes are highlighted in bold text and the same colour as the associated correlation.

|  |  |  | Strength | Clustering coefficient | Closeness |
| --- | --- | --- | --- | --- | --- |
| NAO index | Yearly | ISC | -0.377 ($\chi^2 = 1597$, $p = 0.206$) | 0.996 ($\chi^2 = 1.274$, $p = 0.259$) | **-0.957 ($\chi^2 = 5.269$, $p = 0.022$)** |
| | | RS | Absent ($\chi_{0,1}^2 = 1.688$, $p = 0.097$) | Absent ($\chi_{0,1}^2 = 0.023$, $p = 0.500$) | **Present \*** |
| | Monthly | ISC | -0.976 ($\chi^2 = 0.868$, $p = 0.352$) | 0.075 ($\chi^2 = 0.000$, $p = 0.985$) | -0.486 ($\chi^2 = 0.447$, $p = 0.504$) |
| | | RS | Absent ($\chi_{0,1}^2 = 0.001$, $p = 0.486$) | Absent ($\chi_{0,1}^2 = 0.030$, $p = 0.431$) | **Present ($\chi_{0,1}^2 = 3.448$, $p = 0.032$)** |
| Salmon | Yearly | ISC | **-0.603 ($\chi^2 = 7.195$, $p = 0.007$)** | 0.998 ($\chi^2 = 2.660$, $p = 0.103$) | **0.982 ($\chi^2 = 3.967$, $p = 0.046$)** |
| | | RS | **Present \*** | Absent ($\chi_{0,1}^2 = 0.000$, $p = 0.500$) | **Present \*** |
| | Monthly | ISC | **-0.769 ($\chi^2 = 6.191$, $p = 0.013$)** | -0.967 (NA, model did not converge) | -0.999 ($\chi^2 = 2.005$, $p = 0.157$) |
| | | RS | **Present \*** | Absent ($\chi^2 = 0.035$, $p = 0.983$) | Absent ($\chi_{0,1}^2 = 0.000$, $p = 0.500$) |

Closeness did not vary with NAO index in either sex (Fig. 2c), but it did decrease with increasing salmon numbers in both sexes (Fig. 2f; Table 1; full results in Supplementary Material Tables S5 & S6). Dolphins showed a negative mean-plasticity relationship for the NAO index, with individuals with lower than average mean closeness increasing their closeness with increasing NAO indices and individuals with higher than average closeness decreasing their closeness. This also indicates there was individual variation in plasticity (Table 2). In contrast, there was a positive intercept-slope correlation for closeness in response to salmon abundance (Table 2), indicating that individuals with lower means also decreased the most. The marginal repeatabilities were low for closeness: 0.003 in the NAO index model and 0.002 in the salmon abundance model.

For all social traits there was substantial among-year variation and social traits in consecutive years were positively correlated (year to year correlations: strength; r_NAO_ = 0.452, r_Salmon_ = 0.566; clustering coefficient; r_NAO_ = 0.590, r_Salmon_ = 0.657; and closeness; r_NAO_ =0.283, r_Salmon_ = 0.472), showing that, as expected, adjacent years were more similar than non-adjacent years.

In summary, dolphins had lower closeness scores in years of high salmon abundance, but there were no trait-environment associations at the yearly scale for the other two social traits or for the effect of NAO. For closeness, individuals showed variation in plasticity that was related to their mean behaviour for both NAO and salmon abundance, but for clustering coefficient individuals showed no variation in individual plasticity. Individual dolphins showed variation in their plasticity of strength in response to salmon abundance only, with a great spread of values at high than low values. Clustering coefficient showed consistent differences between individuals across environments, but strength and closeness did not.

### Change with environmental variables at a monthly scale

We analysed how dolphin social phenotypes change in response to environmental variables at the monthly scale with a dataset of 88 unique individuals and 320 measures for all traits. Traits had a mean of 3.64 measures (sd = 2.64) each.

Strength increased with monthly salmon abundance for both sexes (Fig. 3d, Table 1), and there was individual variation in plasticity and a negative intercept-slope correlation, with individuals with lower means increasing more than those with higher means (Table 2). However, strength did not respond to the monthly NAO index (Fig. 3a) and there was no individual variation in plasticity due to the NAO (Table 1, full model results in Tables S7-12 in the supplementary materials). As for the yearly models, the marginal repeatability of strength was low (0.003 in the NAO index model, 0.007 in the salmon abundance model).

**Figure 3.**
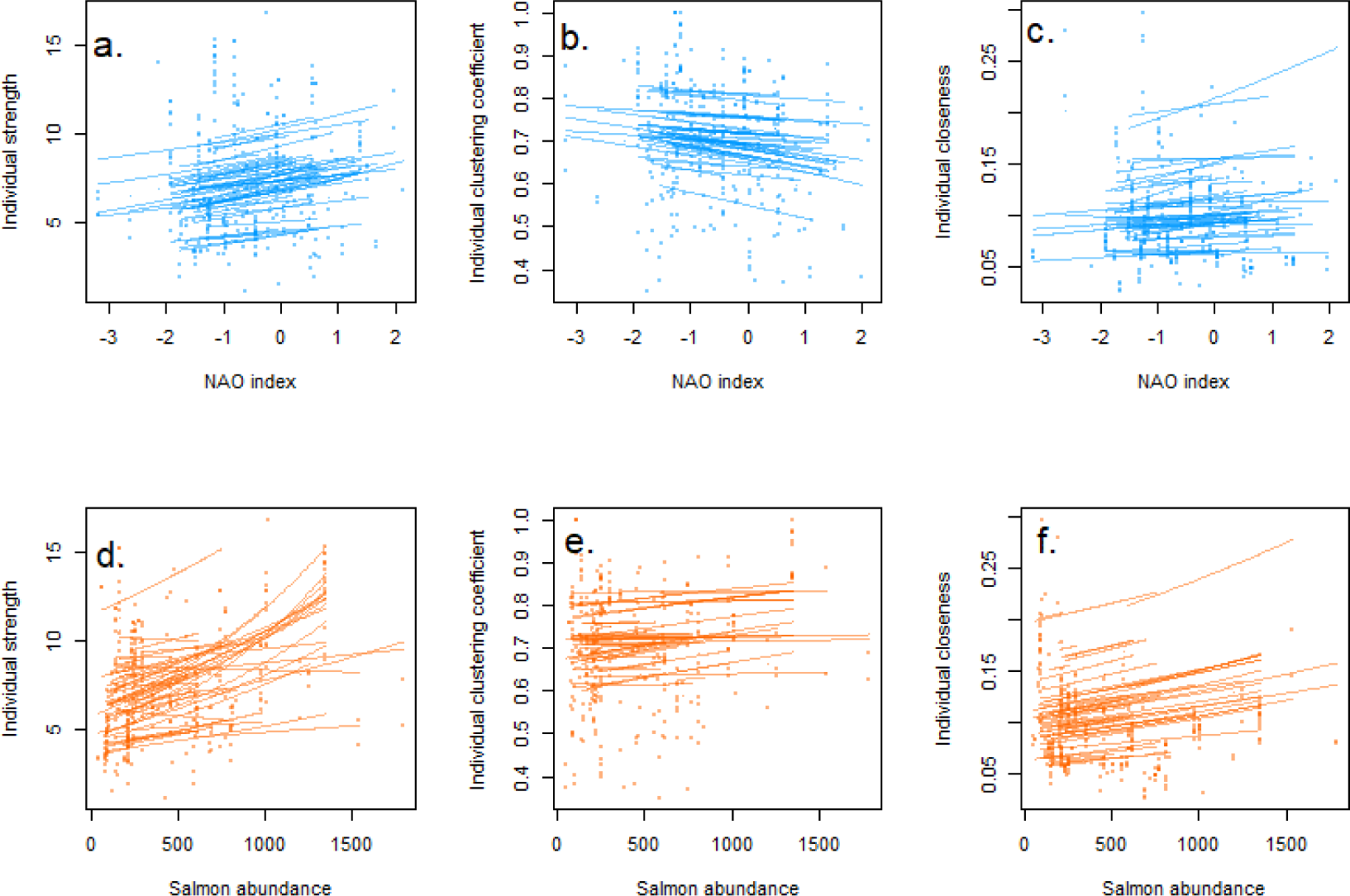
Plots of each of the three social network traits and monthly variation in the North Atlantic Oscillation (NAO) index (a. strength, b. clustering coefficient, c. closeness) and salmon abundances (d. strength, e. clustering coefficient, f. closeness). For each individual dolphin we have predicted its network trait on the observed scale based on the model results, using the “predict” function in R with the individual’s sex, the range of NAO values or salmon counts that individual was exposed to, picking a random year that individual experienced, and the month to June (an arbitrary choice that was approximately in the middle of the calendar year).

Clustering coefficient showed no clear relationships with either the monthly NAO index (Fig. 3b), or the monthly salmon abundance (Fig. 3e and Table 1), and the sexes did not differ in their mean clustering coefficient or how it varied. Individuals showed no mean-plasticity relationship and no individual variation in response to either variable (Table 2, note that the model for clustering coefficient and monthly salmon would not converge with random intercepts and slopes but no correlation between them, so we compared the full model with the model without random slopes using two degrees of freedom). Clustering coefficient was slightly repeatable, with marginal repeatabilites of 0.038 and 0.020 in the NAO index and salmon abundance models respectively.

Both sexes increased their closeness with higher monthly salmon abundance (Fig. 3f), but individuals did not vary in how their closeness changed in response to salmon abundance (Table 2). There was no response to the NAO index (Fig. 3c and Table 1), with no mean-plasticity relationship. However, individuals did vary around the population-level stability in their responses to the NAO index (Table 2). There was limited repeatability of closeness at the monthly scale, with a marginal repeatability of 0.063 in the NAO index model and 0.033 in the salmon abundance model.

There was among-month and among-year variation, but in the monthly models social traits in consecutive years were either negatively or not correlated (year to year correlations: strength: r_NAO_ = −0.016, r_Salmon_ = 0.024; clustering coefficient: r_NAO_ = −0.307, r_Salmon_ = −0.399; closeness: r_NAO_ = −0.144, r_Salmon_ = −0.139).

In summary, months with higher salmon numbers led to higher strength and higher closeness. Meanwhile, the NAO index did not clearly affect any network trait. Individuals differed in how their strength changed in response to salmon abundance, and they also showed variation in responses of closeness to the NAO index, but they did not differ in how their clustering coefficient changed for either variable. Clustering coefficient and closeness were slightly repeatable across environments but strength was not.

## Discussion

We explored whether bottlenose dolphin social behaviour responded to environmental variation. Social behaviour responded to variation in food availability, with a measure of connectedness to the wider network (closeness) decreasing at higher salmon abundances at a yearly scale, and both overall gregariousness (strength) and closeness increasing at higher salmon abundances at a monthly scale. Clustering of the local social environment, clustering coefficient, also showed an increase with monthly salmon abundance but this trend was statistically unclear. In contrast, social behaviours showed no population-level responses to climatic change at either scale. In addition, we found that individuals showed consistent differences in mean clustering coefficient, especially at the yearly scale, but not in individual plasticities, while strength and closeness showed some variation in individual plasticity but limited consistent differences in mean behaviour. This plasticity was often, but not always, negatively associated with mean behaviour, causing among individual differences to typically be greater for low values of salmon abundance and the NAO index.

Months of higher salmon abundance led to increase in all three social behaviours, although this was not statistically clear for clustering coefficient (see Fig. 4 for a comparison of networks between months with low vs. high salmon abundance). This indicates that dolphins were increasing all kinds of social associations in response to increased immediate food availability. Similar results have been found in spotted hyenas (*Crocuta crocuta*) where seasons with high prey availability have denser social networks [77], and in Grant’s gazelles (*Nanger granti*), where increased rainfall (and so food availability) led to higher closeness scores [78]. It is presumed that an increase in social interactions at higher food availability is facilitated by a reduction in the intensity of resource competition [reviewed in:, 6,see also:, 79]. Therefore, in our study system higher rates of social interaction may be beneficial for non-foraging reasons, such as mating. Both mating and calving seasons approximately align with the months of highest salmon abundance in the summer (Fig. S4), supporting this suggestion. Additionally, dolphins could move into areas where food availability is especially high [42,80], causing more individuals to be seen together and therefore inferred social networks to be denser, even if actual rates of social interaction are not changing, or only changing as a byproduct. Finally, it is possible that months with fewer salmon also differ in an unidentified variable which causes dolphins to group less. However, it is unlikely that this unidentified factor is predation threat that changes month to month, as predators are absent in this area [81]. Therefore, a change in social behaviour and/or movement related to the seasonal availability of salmon, perhaps influenced by reproductive behaviour, seems the most likely.

**Figure 4.**
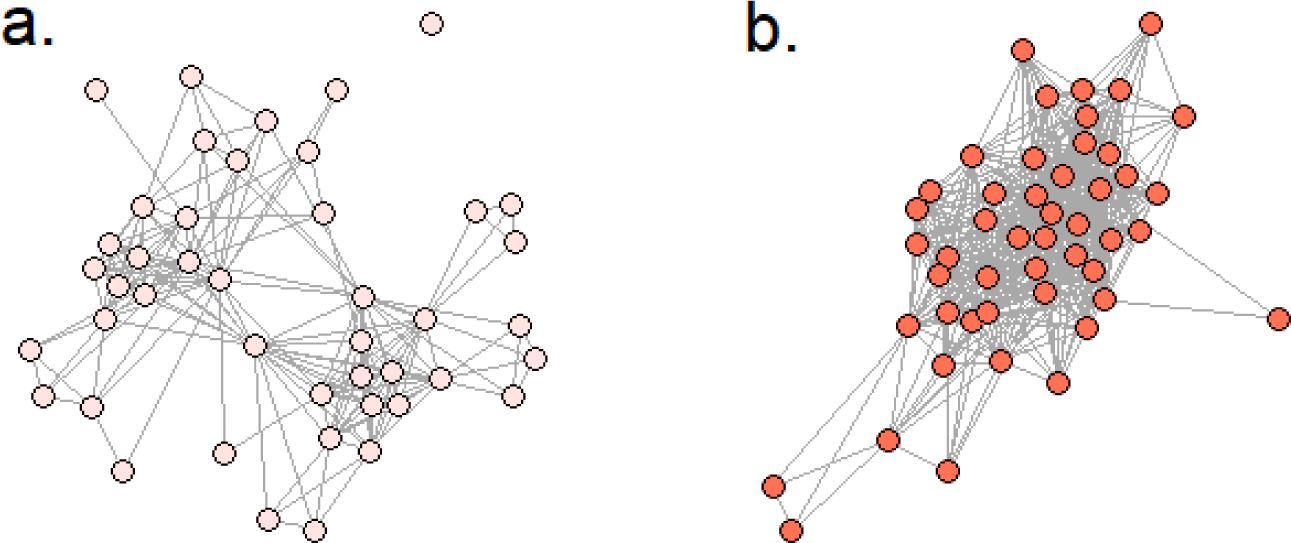
Plots of dolphin social networks in a month of low salmon abundance (May 1990) and a month of high salmon abundance (August 2007). Circles are individual dolphins and grey lines indicate associations i.e., those seen in the same group at least once in that month.

Interestingly, the change in closeness at the yearly scale was in opposite direction to that at the monthly scale, showing a decrease with increased salmon abundance, as well as being a considerably stronger effect (standardised effect size of −0.258 compared to 0.087). A decrease in yearly closeness suggests dolphins are more poorly connected to distant parts of the network when salmon are more abundant. This decrease might be indicative of a reduced need to travel long distances to find food, generating a network with a smaller diameter and so lower individual closeness [see also 82 who demonstrated that more patchy food availability increases network connectedness]. This effect might not be apparent at the monthly scale as the increase in strength in months of high salmon abundance could also lead to higher closeness scores. Whether this is the case or not, the fact that changes in different timescales can be in opposite directions is intriguing and should be kept in mind when attempting to generalise results from one timescale to another.

Despite the changes in response to salmon abundance, we did not see any responses to climatic variation (the NAO index) on a yearly or monthly scale. The NAO index varied considerably at both scales, ranging from −1.15 to 1.08 at the yearly scale and −3.18 to 2.12 at the monthly scale (Fig. S3), hence a lack of necessary variability seems unlikely. Lusseau *et al*. [33] also observed no variation in group size in our study population in response to contemporary variation in the NAO index (they did see an effect at a two-year lag, likely mediated by food availability). The robustness or inflexibility of social behaviour in response to variation in climatic conditions might indicate that the variation in the NAO index is inconsequential and so they have no need to respond to it. Additionally or alternatively, dolphins may change other phenotypes, such as foraging behaviour or metabolism, to cope with this stressor [31], leaving social behaviour unchanged. Finally, local conditions might be more relevant to dolphin behaviour, as opposed to the regional conditions summarised by the NAO index. For example, movements of bottlenose dolphins depend on tidal currents and fronts [83], and changes to these might be important for their social behaviour.

Alongside the plasticity at the population level in strength and closeness, these traits also showed individual variation in plasticity (although strength only showed this for salmon abundance, and closeness only at the yearly level). Therefore, even if the population as a whole showed no overall change for some trait-environmental variable combinations, some individuals might still show an increase in their overall gregariousness and/or connectedness to wider parts of the network, while others a decrease. Variation in individual plasticity leads to environment-dependent repeatability (and possibly heritability), can dampen population responses to environmental variability, and enhance population persistence [18]. For example, due to the negative intercept-slope correlation, individual strength shows the most among-individual variation at low salmon abundances (the approximate marginal repeatability of strength at the monthly scale two standard deviations below the mean salmon abundance was 0.124, compared to 0.02 at the mean). If strength is linked to foraging strategy, for instance if individuals with more social connections have access to more information about prey availability [82], a wider range of social phenotypes could increase the possibility that at least some individuals are successful despite low food availability. Determining the genetic basis and importance of early life conditions for the development of these different responses [12] and how this variation impacts population dynamics [84] is key. Models projecting how these traits in dolphin populations will change in the future should account for both among-individual variation in mean behaviour and in behavioural plasticity. Additionally, Kebke *et al*. [31] suggest that cetacean ranging and foraging behaviour may well be under selection for increased plasticity as environments change, and so estimating selection on both means and plasticities of behaviours is a logical and important next step.

In contrast, clustering coefficient showed no individual variation in plasticity at any timescale-environmental variable combination. Clustering coefficient at the yearly scale was the only trait with more than slight repeatability, indicating some individual consistency [see also 85 who found consistency over lifetime in the social behaviours of Indo-Pacific bottlenose dolphins, *T. aduncus*]. Therefore, plasticity might be more limited with individuals keeping the same pattern of local connections across environmental conditions. As clustering coefficient depends on the frequency of connections among triads, an individual’s trait value cannot change without impacting the trait value of others. This interdependence may then constrain the degree of plasticity possible at the individual level. There is no evidence for male alliances in this population [86], and so determining what these clusters of individuals represent and why they might be so stable would be useful.

In conclusion, we found that, at the population level, individual dolphin social behaviour is more responsive to variation in food availability than climatic variation, with this being particularly apparent at the monthly scale. We observed that individuals increased overall gregariousness and connectedness to wider parts of the network in months of higher salmon abundance. In contrast, dolphins decreased their connectedness to wider parts of the network in years of high salmon availability. Traits that tended to show higher repeatability tended to show limited individual variation in plasticity, although there was considerable variation in this trend. As such, whether individual heterogeneity in both mean and plasticity in behaviour needs to be accounted for when predicting species responses to environmental change might have to be considered on a case-by-case basis, and individual plasticities as well as means may be targets of selection and hence evolvable.

## Acknowledgements

We are indebted to Paul Thompson for conception and development of this long-term individual based research study, and for advice during this work. We are especially grateful to our colleagues at the Lighthouse Field Station, past and present, who have contributed to and advanced this long-term study. Matthew Silk, Dan Blumstein, and two anonymous reviewers made numerous useful comments and suggestions on an earlier draft. This long-term study has depended upon a number of funders including University of Aberdeen, NatureScot, Beatrice Offshore Windfarm Ltd., Moray Offshore Renewables Ltd., Marine Scotland, the Crown Estate, Highlands and Islands Enterprise, the BES, ASAB, Greenpeace Environmental Trust, Whale and Dolphin Conservation, Talisman Energy (UK) Ltd., Department of Energy and Climate Change, Chevron and Natural Environment Research. Survey work was conducted under NatureScot Animal Scientific Licences. We have no conflicts of interest.

## Statement of authorship

DF conceived the idea for the manuscript, BC provided data, DF ran analyses with guidance from BC, DF wrote first draft of the manuscript and both authors contributed to revisions.

## Data accessibility statement

Data and R code used for the analysis are available at [76].

