## Supplementary materials for "Dolphin social phenotypes vary in response to food availability but not the North Atlantic Oscillation index"

### Supplementary Figures

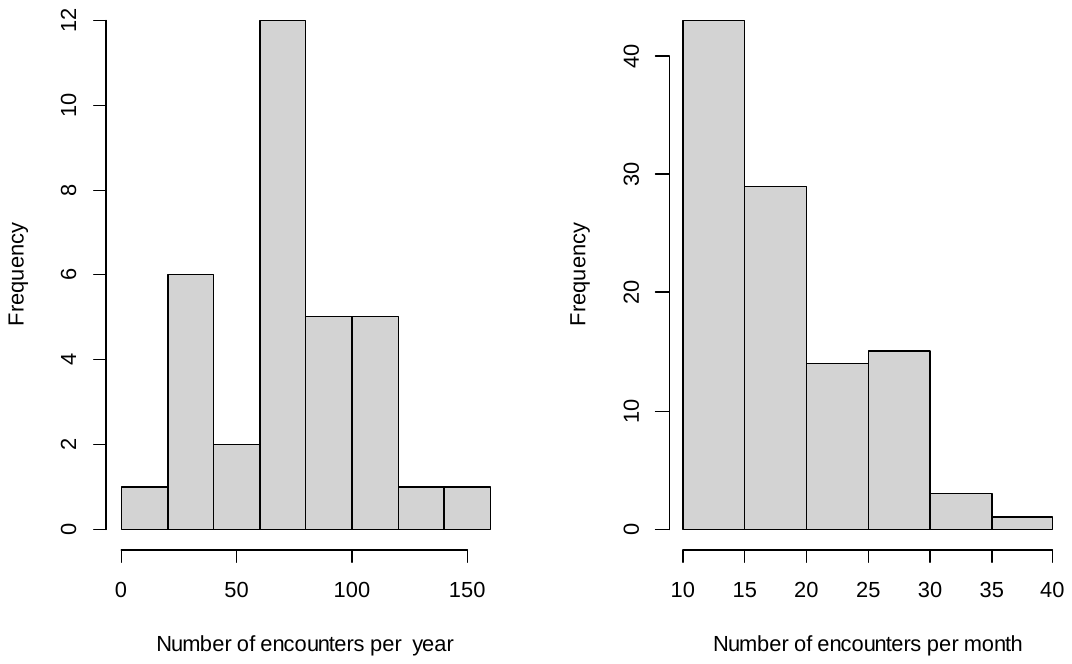

**Figure S1**. Histograms of the number of unique encounters with dolphin groups that were used to build the Yearly (left) and Monthly (right) networks.

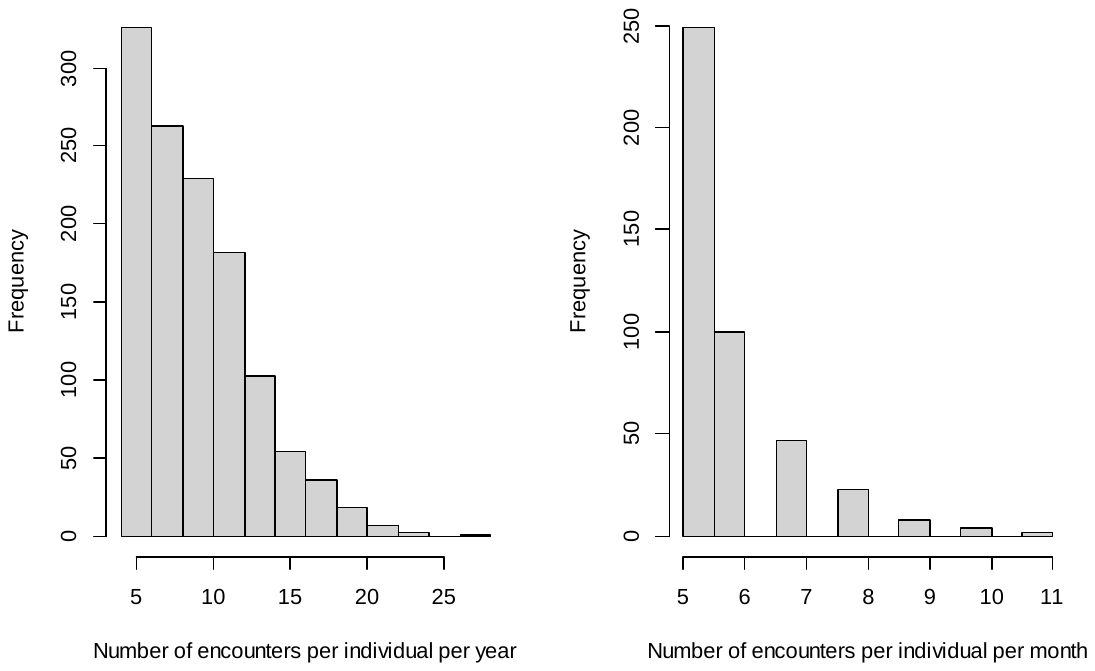

**Figure S2**. Histograms of the number of encounters that each individual was seen in per Year (left) and per Month (right) during the study period.

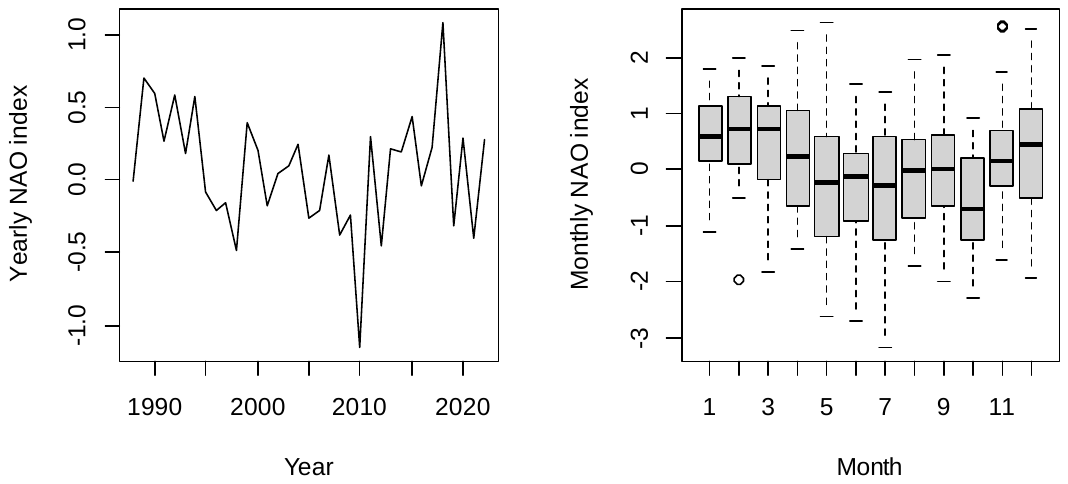

**Figure S3**. Plots of the yearly NAO index (left; calculated as the average of the 12 months that year) and the monthly NAO index per month over the study period.

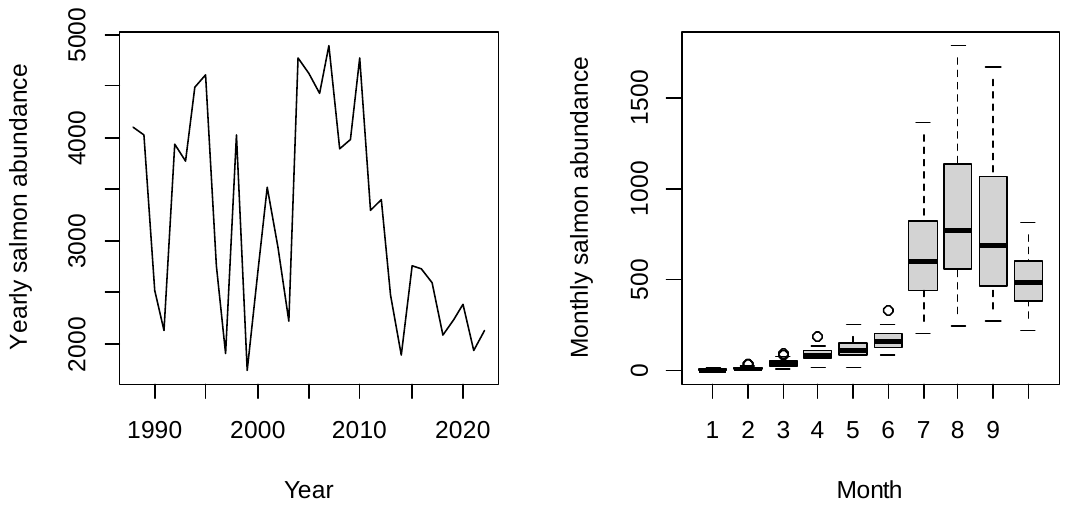

**Figure S4**. Plots of the yearly salmon abundance (summed across all months) and the monthly salmon catch per month over the study period (no salmon were caught in November or December).

### Supplementary Tables

Tables S1-12. Full results from the models described in the main text. In the top half of the table are shown the fixed effects, with effect estimate, standard error, the χ^2^ statistic, and the p-value. Degrees of freedom is one in all cases. For sex males are the default category and females the contrast. In the bottom half of the table are shown the random terms (either effects or correlations) and their associated variance component or correlation coefficient. Tables S1-6 are for models at the yearly scale, Tables S7-12 for models at the monthly scale.

**Table S1**. Model with yearly strength as the response variable and the North Atlantic Oscillation (NAO) index as the environmental variable in the predictors.

| Variable | Estimate | Standard Error | Chi-squared | P value |
| --- | --- | --- | --- | --- |
| Intercept | 1.614 | 0.101 | 257.097 | 0.000 |
| NAOI (scaled) | -0.035 | 0.066 | 0.280 | 0.597 |
| Sex (male contrast) | -0.044 | 0.046 | 0.916 | 0.339 |
| NAOI : Sex | -0.002 | 0.021 | 0.005 | 0.942 |
|  |  | Random effect/Correlation | Variance |  |
|  |  | Individual Intercepts | 0.044 |  |
|  |  | Individual Slopes | 0.002 |  |
|  |  | Intercept - slope correlation | -0.377 |  |
|  |  | Year | 0.102 |  |
|  |  | Year autocorrelation | 0.452 |  |

**Table S2**. Model with yearly strength as the response variable and salmon abundance as the environmental variable in the predictors.

| Variable | Estimate | Standard Error | Chi-squared | P value |
| --- | --- | --- | --- | --- |
| Intercept | 1.601 | 0.117 | 187.721 | 0.000 |
| Wild salmon numbers (scaled) | 0.108 | 0.069 | 2.422 | 0.120 |
| Sex (male contrast) | -0.049 | 0.046 | 1.157 | 0.282 |
| Wild salmon : Sex | -0.009 | 0.026 | 0.126 | 0.723 |
|  |  | Random effect/Correlation | Variance |  |
|  |  | Individual Intercepts | 0.042 |  |
|  |  | Individual Slopes | 0.004 |  |
|  |  | Intercept - slope correlation | -0.603 |  |
|  |  | Year | 0.110 |  |
|  |  | Year autocorrelation | 0.566 |  |

**Table S3**. Model with yearly clustering coefficient as the response variable and the North Atlantic Oscillation (NAO) index as the environmental variable in the predictors.

| Variable | Estimate | Standard Error | Chi-squared | P value |
| --- | --- | --- | --- | --- |
| Intercept | 0.780 | 0.169 | 21.216 | 0.000 |
| NAOI (scaled) | -0.102 | 0.081 | 1.595 | 0.207 |
| Sex (male contrast) | 0.074 | 0.056 | 1.706 | 0.192 |
| NAOI : Sex | 0.031 | 0.035 | 0.793 | 0.373 |
|  |  | Random effect/Correlation | Variance |  |
|  |  | Individual Intercepts | 0.050 |  |
|  |  | Individual Slopes | 0.001 |  |
|  |  | Intercept - slope correlation | 0.996 |  |
|  |  | Year | 0.226 |  |
|  |  | Year autocorrelation | 0.590 |  |

**Table S4**. Model with yearly clustering coefficient as the response variable and salmon abundance as the environmental variable in the predictors.

| Variable | Estimate | Standard Error | Chi-squared | P value |
| --- | --- | --- | --- | --- |
| Intercept | 0.768 | 0.191 | 21.215 | 0.000 |
| Wild salmon (scaled) | -0.027 | 0.091 | 1.595 | 0.207 |
| Sex (male contrast) | 0.074 | 0.057 | 1.706 | 0.192 |
| Wild salmon : Sex | -0.070 | 0.038 | 0.794 | 0.373 |
|  |  | Random effect/Correlation | Variance |  |
|  |  | Individual Intercepts | 0.053 |  |
|  |  | Individual Slopes | 0.001 |  |
|  |  | Intercept - slope correlation | 0.998 |  |
|  |  | Year | 0.240 |  |
|  |  | Year autocorrelation | 0.657 |  |

**Table S5**. Model with yearly closeness as the response variable and the North Atlantic Oscillation (NAO) index as the environmental variable in the predictors.

| Variable | Estimate | Standard Error | Chi-squared | P value |
| --- | --- | --- | --- | --- |
| Intercept | -2.357 | 0.119 | 394.951 | 0.000 |
| NAOI (scaled) | -0.115 | 0.102 | 1.275 | 0.259 |
| Sex (male contrast) | 0.006 | 0.009 | 0.405 | 0.525 |
| NAOI : Sex | 0.006 | 0.008 | 0.458 | 0.499 |
|  |  | Random effect/Correlation | Variance |  |
|  |  | Individual Intercepts | 0.001 |  |
|  |  | Individual Slopes | 0.000 |  |
|  |  | Intercept - slope correlation | -0.957 |  |
|  |  | Year | 0.246 |  |
|  |  | Year autocorrelation | 0.283 |  |

**Table S6**. Model with yearly closeness as the response variable and salmon abundance as the environmental variable in the predictors.

| Variable | Estimate | Standard Error | Chi-squared | P value |
| --- | --- | --- | --- | --- |
| Intercept | -2.354 | 0.143 | 270.986 | 0.000 |
| Wild salmon (scaled) | -0.258 | 0.094 | 7.489 | 0.006 |
| Sex (male contrast) | 0.004 | 0.009 | 0.189 | 0.664 |
| Wild salmon : Sex | -0.002 | 0.008 | 0.091 | 0.763 |
|  |  | Random effect/Correlation | Variance |  |
|  |  | Individual Intercepts | 0.001 |  |
|  |  | Individual Slopes | 0.000 |  |
|  |  | Intercept - slope correlation | 0.982 |  |
|  |  | Year | 0.235 |  |
|  |  | Year autocorrelation | 0.472 |  |

**Table S7**. Model with monthly strength as the response variable and the North Atlantic Oscillation (NAO) index as the environmental variable in the predictors.

| Variable | Estimate | Standard Error | Chi-squared | P value |
| --- | --- | --- | --- | --- |
| Intercept | 1.962 | 0.102 | 369.864 | 0.000 |
| NAOI (scaled) | 0.053 | 0.034 | 2.437 | 0.119 |
| Sex (male contrast) | -0.043 | 0.046 | 0.862 | 0.353 |
| NAOI : Sex | 0.039 | 0.037 | 1.105 | 0.293 |
|  |  | Random effect/Correlation | Variance |  |
|  |  | Individual Intercepts | 0.013 |  |
|  |  | Individual Slopes | 0.001 |  |
|  |  | Intercept - slope correlation | -0.976 |  |
|  |  | Month | 0.025 |  |
|  |  | Year | 0.086 |  |
|  |  | Year autocorrelation | -0.016 |  |

**Table S8**. Model with monthly strength as the response variable and salmon abundance as the environmental variable in the predictors.

| Variable | Estimate | Standard Error | Chi-squared | P value |
| --- | --- | --- | --- | --- |
| Intercept | 1.963 | 0.070 | 787.986 | 0.000 |
| Wild salmon (scaled) | 0.137 | 0.038 | 12.755 | 0.000 |
| Sex (male contrast) | -0.044 | 0.050 | 0.772 | 0.380 |
| Wild salmon : Sex | -0.011 | 0.045 | 0.065 | 0.799 |
|  |  | Random effect/Correlation | Variance |  |
|  |  | Individual Intercepts | 0.017 |  |
|  |  | Individual Slopes | 0.015 |  |
|  |  | Intercept - slope correlation | -0.769 |  |
|  |  | Month | 0.001 |  |
|  |  | Year | 0.066 |  |
|  |  | Year autocorrelation | 0.024 |  |

**Table S9**. Model with monthly clustering coefficient as the response variable and the North Atlantic Oscillation (NAO) index as the environmental variable in the predictors.

| Variable | Estimate | Standard Error | Chi-squared | P value |
| --- | --- | --- | --- | --- |
| Intercept | 0.805 | 0.073 | 121.200 | 0.000 |
| NAOI (scaled) | -0.102 | 0.056 | 3.304 | 0.069 |
| Sex (male contrast) | 0.107 | 0.070 | 2.352 | 0.125 |
| NAOI : Sex | 0.057 | 0.069 | 0.678 | 0.410 |
|  |  | Random effect/Correlation | Variance |  |
|  |  | Individual Intercepts | 0.003 |  |
|  |  | Individual Slopes | 0.002 |  |
|  |  | Intercept - slope correlation | 0.075 |  |
|  |  | Month | 0.000 |  |
|  |  | Year | 0.095 |  |
|  |  | Year autocorrelation | -0.307 |  |

**Table S10**. Model with monthly clustering coefficient as the response variable and salmon abundance as the environmental variable in the predictors.

| Variable | Estimate | Standard Error | Chi-squared | P value |
| --- | --- | --- | --- | --- |
| Intercept | 0.793 | 0.071 | 123.094 | 0.000 |
| Wild salmon (scaled) | 0.116 | 0.062 | 3.569 | 0.059 |
| Sex (male contrast) | 0.100 | 0.070 | 2.071 | 0.150 |
| Wild salmon : Sex | -0.105 | 0.074 | 2.011 | 0.156 |
|  |  | Random effect/Correlation | Variance |  |
|  |  | Individual Intercepts | 0.003 |  |
|  |  | Individual Slopes | 0.000 |  |
|  |  | Intercept - slope correlation | -0.967 |  |
|  |  | Month | 0.000 |  |
|  |  | Year | 0.111 |  |
|  |  | Year autocorrelation | -0.399 |  |

**Table S11**. Model with monthly closeness as the response variable and the North Atlantic Oscillation (NAO) index as the environmental variable in the predictors.

| Variable | Estimate | Standard Error | Chi-squared | P value |
| --- | --- | --- | --- | --- |
| Intercept | -2.417 | 0.113 | 455.958 | 0.000 |
| NAOI (scaled) | 0.023 | 0.033 | 0.464 | 0.496 |
| Sex (male contrast) | 0.009 | 0.031 | 0.087 | 0.768 |
| NAOI : Sex | 0.031 | 0.037 | 0.713 | 0.398 |
|  |  | Random effect/Correlation | Variance |  |
|  |  | Individual Intercepts | 0.002 |  |
|  |  | Individual Slopes | 0.005 |  |
|  |  | Intercept - slope correlation | -0.486 |  |
|  |  | Month | 0.044 |  |
|  |  | Year | 0.094 |  |
|  |  | Year autocorrelation | -0.144 |  |

**Table S12**. Model with monthly clustering coefficient as the response variable and salmon abundance as the environmental variable in the predictors.

| Variable | Estimate | Standard Error | Chi-squared | P value |
| --- | --- | --- | --- | --- |
| Intercept | -2.418 | 0.143 | 287.409 | 0.000 |
| Wild salmon (scaled) | 0.087 | 0.040 | 4.714 | 0.030 |
| Sex (male contrast) | 0.006 | 0.033 | 0.030 | 0.863 |
| Wild salmon : Sex | 0.009 | 0.030 | 0.081 | 0.777 |
|  |  | Random effect/Correlation | Variance |  |
|  |  | Individual Intercepts | 0.003 |  |
|  |  | Individual Slopes | 0.001 |  |
|  |  | Intercept - slope correlation | -0.999 |  |
|  |  | Month | 0.081 |  |
|  |  | Year | 0.093 |  |
|  |  | Year autocorrelation | -0.139 |  |

Analysis with minimum three observations

In the main text we show results of analyses where dolphins are only included if they have a minimum of five observations per year or per monthly for those respective analyses. However, we initially performed the analysis with a lower threshold of three observations. We show these results here.

For the yearly analysis, we removed the lone individual with a strength of zero, giving 1167 observations of 157 unique individuals. For the monthly analysis, we again removed a single individual with a strength of zero, giving 1357 observations of 138 unique individuals. Results of each model are given in tables S13-24. In the top half of the table are shown the fixed effects, with effect estimate, standard error, the χ2 statistic, and the p-value. Degrees of freedom is one in all cases. For sex males are the default category and females the contrast. In the bottom half of the table are shown the random terms (either effects or correlations) and their associated variance component or correlation coefficient. Tables S13-18 are for models at the yearly scale, Tables S19-24 for models at the monthly scale.

Table S13. Model with yearly strength as the response variable and the North Atlantic Oscillation (NAO) index as the environmental variable in the predictors.

| Variable | Estimate | Standard Error | Chi-squared | P value |
| --- | --- | --- | --- | --- |
| Intercept | 1.606 | 0.092 | 301.454 | 0.000 |
| NAOI (scaled) | -0.049 | 0.065 | 0.565 | 0.452 |
| Sex (male contrast) | -0.026 | 0.049 | 0.272 | 0.602 |
| NAOI : Sex | -0.006 | 0.023 | 0.061 | 0.805 |
|  |  | Random effect/Correlation | Variance |  |
|  |  | Individual Intercepts | 0.061 |  |
|  |  | Individual Slopes | 0.003 |  |
|  |  | Intercept - slope correlation | -0.662 |  |
|  |  | Year | 0.112 |  |
|  |  | Year autocorrelation | 0.333 |  |

Table S14. Model with yearly strength as the response variable and salmon abundance as the environmental variable in the predictors.

| Variable | Estimate | Standard Error | Chi-squared | P value |
| --- | --- | --- | --- | --- |
| Intercept | 1.600 | 0.112 | 203.006 | 0.000 |
| Wild salmon numbers (scaled) | 0.142 | 0.071 | 3.975 | 0.046 |
| Sex (male contrast) | -0.026 | 0.049 | 0.288 | 0.591 |
| Wild salmon : Sex | 0.003 | 0.028 | 0.008 | 0.927 |
|  |  | Random effect/Correlation | Variance |  |
|  |  | Individual Intercepts | 0.057 |  |
|  |  | Individual Slopes | 0.007 |  |
|  |  | Intercept - slope correlation | -0.537 |  |
|  |  | Year | 0.119 |  |
|  |  | Year autocorrelation | 0.514 |  |

Table S15. Model with yearly clustering coefficient as the response variable and the North Atlantic Oscillation (NAO) index as the environmental variable in the predictors.

| Variable | Estimate | Standard Error | Chi-squared | P value |
| --- | --- | --- | --- | --- |
| Intercept | 0.870 | 0.144 | 36.260 | 0.000 |
| NAOI (scaled) | -0.107 | 0.077 | 1.904 | 0.168 |
| Sex (male contrast) | 0.032 | 0.052 | 0.384 | 0.535 |
| NAOI : Sex | 0.012 | 0.035 | 0.113 | 0.737 |
|  |  | Random effect/Correlation | Variance |  |
|  |  | Individual Intercepts | 0.046 |  |
|  |  | Individual Slopes | 0.000 |  |
|  |  | Intercept - slope correlation | 0.998 |  |
|  |  | Year | 0.205 |  |
|  |  | Year autocorrelation | 0.518 |  |

Table S16. Model with yearly clustering coefficient as the response variable and salmon abundance as the environmental variable in the predictors.

| Variable | Estimate | Standard Error | Chi-squared | P value |
| --- | --- | --- | --- | --- |
| Intercept | 0.866 | 0.158 | 36.260 | 0.000 |
| Wild salmon (scaled) | 0.036 | 0.086 | 1.904 | 0.168 |
| Sex (male contrast) | 0.036 | 0.053 | 0.384 | 0.535 |
| Wild salmon : Sex | 0.003 | 0.042 | 0.113 | 0.737 |
|  |  | Random effect/Correlation | Variance |  |
|  |  | Individual Intercepts | 0.046 |  |
|  |  | Individual Slopes | 0.009 |  |
|  |  | Intercept - slope correlation | 0.339 |  |
|  |  | Year | 0.226 |  |
|  |  | Year autocorrelation | 0.558 |  |

Table S17. Model with yearly closeness as the response variable and the North Atlantic Oscillation (NAO) index as the environmental variable in the predictors.

| Variable | Estimate | Standard Error | Chi-squared | P value |
| --- | --- | --- | --- | --- |
| Intercept | -2.589 | 0.157 | 272.201 | 0.000 |
| NAOI (scaled) | -0.273 | 0.186 | 2.166 | 0.141 |
| Sex (male contrast) | 0.139 | 0.108 | 1.655 | 0.198 |
| NAOI : Sex | 0.081 | 0.056 | 2.094 | 0.148 |
|  |  | Random effect/Correlation | Variance |  |
|  |  | Individual Intercepts | 0.385 |  |
|  |  | Individual Slopes | 0.094 |  |
|  |  | Intercept - slope correlation | 1.000 |  |
|  |  | Year | 0.762 |  |
|  |  | Year autocorrelation | -0.168 |  |

Table S18. Model with yearly closeness as the response variable and salmon abundance as the environmental variable in the predictors.

| Variable | Estimate | Standard Error | Chi-squared | P value |
| --- | --- | --- | --- | --- |
| Intercept | -2.539 | 0.147 | 300.089 | 0.000 |
| Wild salmon (scaled) | -0.201 | 0.150 | 1.789 | 0.181 |
| Sex (male contrast) | 0.153 | 0.076 | 3.987 | 0.046 |
| Wild salmon : Sex | 0.173 | 0.076 | 5.220 | 0.022 |
|  |  | Random effect/Correlation | Variance |  |
|  |  | Individual Intercepts | 0.158 |  |
|  |  | Individual Slopes | 0.150 |  |
|  |  | Intercept - slope correlation | 0.734 |  |
|  |  | Year | 0.674 |  |
|  |  | Year autocorrelation | -0.097 |  |

Table S19. Model with monthly strength as the response variable and the North Atlantic Oscillation (NAO) index as the environmental variable in the predictors.

| Variable | Estimate | Standard Error | Chi-squared | P value |
| --- | --- | --- | --- | --- |
| Intercept | 1.810 | 0.121 | 224.998 | 0.000 |
| NAOI (scaled) | 0.030 | 0.023 | 1.746 | 0.186 |
| Sex (male contrast) | 0.008 | 0.025 | 0.098 | 0.754 |
| NAOI : Sex | 0.008 | 0.025 | 0.093 | 0.760 |
|  |  | Random effect/Correlation | Variance |  |
|  |  | Individual Intercepts | 0.000 |  |
|  |  | Individual Slopes | 0.000 |  |
|  |  | Intercept - slope correlation | -0.679 |  |
|  |  | Month | 0.034 |  |
|  |  | Year | 0.078 |  |
|  |  | Year autocorrelation | 0.433 |  |

Table S20. Model with monthly strength as the response variable and salmon abundance as the environmental variable in the predictors.

| Variable | Estimate | Standard Error | Chi-squared | P value |
| --- | --- | --- | --- | --- |
| Intercept | 1.839 | 0.115 | 256.604 | 0.000 |
| Wild salmon (scaled) | 0.037 | 0.029 | 1.655 | 0.198 |
| Sex (male contrast) | -0.006 | 0.055 | 0.011 | 0.917 |
| Wild salmon : Sex | 0.028 | 0.028 | 0.941 | 0.332 |
|  |  | Random effect/Correlation | Variance |  |
|  |  | Individual Intercepts | 0.066 |  |
|  |  | Individual Slopes | 0.010 |  |
|  |  | Intercept - slope correlation | -0.981 |  |
|  |  | Month | 0.020 |  |
|  |  | Year | 0.079 |  |
|  |  | Year autocorrelation | 0.424 |  |

Table S21. Model with monthly clustering coefficient as the response variable and the North Atlantic Oscillation (NAO) index as the environmental variable in the predictors.

| Variable | Estimate | Standard Error | Chi-squared | P value |
| --- | --- | --- | --- | --- |
| Intercept | 1.046 | 0.107 | 95.197 | 0.000 |
| NAOI (scaled) | -0.068 | 0.040 | 2.932 | 0.087 |
| Sex (male contrast) | 0.040 | 0.053 | 0.560 | 0.454 |
| NAOI : Sex | 0.088 | 0.045 | 3.866 | 0.049 |
|  |  | Random effect/Correlation | Variance |  |
|  |  | Individual Intercepts | 0.020 |  |
|  |  | Individual Slopes | 0.002 |  |
|  |  | Intercept - slope correlation | -0.506 |  |
|  |  | Month | 0.014 |  |
|  |  | Year | 0.064 |  |
|  |  | Year autocorrelation | 0.486 |  |

Table S22. Model with monthly clustering coefficient as the response variable and salmon abundance as the environmental variable in the predictors.

| Variable | Estimate | Standard Error | Chi-squared | P value |
| --- | --- | --- | --- | --- |
| Intercept | 1.039 | 0.105 | 98.584 | 0.000 |
| Wild salmon (scaled) | 0.180 | 0.052 | 12.118 | 0.000 |
| Sex (male contrast) | 0.044 | 0.053 | 0.686 | 0.407 |
| Wild salmon : Sex | -0.076 | 0.048 | 2.588 | 0.108 |
|  |  | Random effect/Correlation | Variance |  |
|  |  | Individual Intercepts | 0.020 |  |
|  |  | Individual Slopes | 0.000 |  |
|  |  | Intercept - slope correlation | 0.989 |  |
|  |  | Month | 0.020 |  |
|  |  | Year | 0.061 |  |
|  |  | Year autocorrelation | 0.368 |  |

Table S23. Model with monthly closeness as the response variable and the North Atlantic Oscillation (NAO) index as the environmental variable in the predictors.

| Variable | Estimate | Standard Error | Chi-squared | P value |
| --- | --- | --- | --- | --- |
| Intercept | -2.553 | 0.146 | 303.797 | 0.000 |
| NAOI (scaled) | 0.040 | 0.041 | 0.927 | 0.336 |
| Sex (male contrast) | -0.034 | 0.052 | 0.422 | 0.516 |
| NAOI : Sex | 0.039 | 0.047 | 0.679 | 0.410 |
|  |  | Random effect/Correlation | Variance |  |
|  |  | Individual Intercepts | 0.040 |  |
|  |  | Individual Slopes | 0.020 |  |
|  |  | Intercept - slope correlation | 0.126 |  |
|  |  | Month | 0.069 |  |
|  |  | Year | 0.108 |  |
|  |  | Year autocorrelation | 0.173 |  |

Table S24. Model with monthly closeness as the response variable and salmon abundance as the environmental variable in the predictors.

| Variable | Estimate | Standard Error | Chi-squared | P value |
| --- | --- | --- | --- | --- |
| Intercept | -2.577 | 0.127 | 408.704 | 0.000 |
| Wild salmon (scaled) | -0.115 | 0.057 | 4.049 | 0.044 |
| Sex (male contrast) | -0.003 | 0.054 | 0.003 | 0.960 |
| Wild salmon : Sex | 0.082 | 0.054 | 2.323 | 0.127 |
|  |  | Random effect/Correlation | Variance |  |
|  |  | Individual Intercepts | 0.040 |  |
|  |  | Individual Slopes | 0.032 |  |
|  |  | Intercept - slope correlation | -0.388 |  |
|  |  | Month | 0.043 |  |
|  |  | Year | 0.096 |  |
|  |  | Year autocorrelation | 0.226 |  |
